# BiXformer: A Bidirectional Cross Attention Transformer for Disentangling Inter-Regional Neural Dynamics

**DOI:** 10.64898/2026.06.05.730511

**Authors:** Omar El Sayed, Yujin Han, Tudor Dragoi, Michael N. Economo, Brian DePasquale

## Abstract

Advances in high-throughput neural recording technologies enable simultaneous measurement of activity across multiple brain regions in behaving animals, producing datasets of unprecedented scale and richness. Interpreting these data remains challenging due to the bidirectional and temporally offset nature of inter-regional communication, where feedforward and feedback signals are superimposed within neural populations. We introduce BiXformer, a bidirectional cross-attention transformer that disentangles these interactions by decomposing inter-regional communication into causal and acausal streams using directionally masked attention. By enforcing temporal constraints within attention heads, BiXformer recovers low-dimensional, directed latent dynamics and estimates communication delays without relying on linearity or stationarity assumptions. We validate the model on synthetic datasets with known ground-truth delays, demonstrating accurate recovery of both latent structure and inter-regional timing. Applied to simultaneous neural-behavioral recordings and multi-region neural recordings during a movement task, BiXformer reveals interpretable, temporally structured components consistent with the coexistence of sensory feedback and motor-related signals. These results establish BiXformer as a flexible framework for uncovering dynamic, directed communication in complex neural circuits.

## 1 Introduction

Advances in recording techniques enable simultaneous measurement of neural activity across multiple regions in a behaving animal, producing datasets of unprecedented scale and richness [1–5]. The same scale that renders these datasets powerful also makes them remarkably difficult to interpret. A key part of this difficulty lies in the brain’s architecture: regions underlying different processes, such as cognition [6, 7], sensation [8–11], and movement [2, 12], are densely interconnected, with information relayed bi-directionally and concurrently through many pathways and across multiple timescales. A central challenge in systems neuroscience is characterizing the information flow between brain regions – specifically, the content, timing, and directionality of inter-regional communication.

This challenge is acute in sensorimotor circuits [13, 14] where neural responses often integrate signals that both drive and reflect movement. For instance, in a rhythmic licking task, neurons in the primary motor cortex can exhibit both feedforward motor commands that drive movement and feedback signals reflecting proprioceptive input arising from movement [15, 16]. Crucially, these signals do not arrive simultaneously; communication between populations is subject to conduction delays that reflect the underlying anatomical organization and physiological properties of neurons [17]. The mixing of these temporally offset signals within a region’s neural population creates a fundamental confound correlational methods cannot resolve. Without explicitly disentangling feedforward from feedback interactions, the direction of information flow – between neural populations or between neural activity and behavior – remains ambiguous.

Addressing this central challenge has motivated a growing body of computational approaches aimed at modeling inter-regional interactions [18–31]. While these approaches have made significant progress, particularly in identifying smooth, low-dimensional latent dynamics across neural populations, they often rely on linearity or stationarity assumptions that ultimately constrain them to a fixed mode of communication, leaving the potential to consider broader forms of communication between regions unexplored. Neural circuits are inherently nonlinear [27] and non-stationary, re-configuring their dynamics across timescales and behavioral contexts [32, 33]. An ideal model of inter-regional communication should be capable of capturing nonlinear interactions between populations, time-dependent changes in communication, and temporal delays inherent to inter-regional signaling.

Transformers offer a principled framework for satisfying these criteria. While their ability to capture temporally-extended, time-dependent interactions in high dimensional sequential data makes them particularly well-suited for modeling neural dynamics, they have largely been used as predictive models rather than tools for mechanistic interpretation.

To address this need, we introduce the *Bidirectional Cross Attention Transformer* (BiXformer), a novel architecture designed to disentangle dynamic and directed information flow across neural populations. BiXformer decomposes inter-regional communication into two parallel streams — a causal stream and an acausal stream — by leveraging masked cross-attention heads to explicitly disentangle feedforward and feedback interactions between populations. In this work, we demonstrate the model’s ability to recover these temporally-asymmetric signals in synthetic data and show that it yields interpretable, directed latent dynamics when applied to real simultaneous neural-behavioral and multi-region recordings from mice performing a controlled motor task.

Our contributions can be summarized as follows:

- **We introduce BiXformer**, a transformer architecture that decomposes inter-regional communication into causal and acausal streams via masked cross-attention.
- **We validate BiXformer on synthetic data**, demonstrating that BiXformer accurately recovers causal and acausal latents and their associated inter-regional delays.
- **We apply BiXformer to simultaneous neural-behavioral recordings** from the medulla (Md) during a rhythmic licking task, revealing interpretable latent components consistent with coexisting sensory and motor signals.
- **We use BiXformer to uncover interactions between primary motor cortex (M1) and primary somatosensory cortex (S1)** during a delayed reward licking task, confirming existing hypotheses about sensorimotor communication while shedding new light on the nature of S1-M1 communication.

## 2 Related Work

### 2.1 Latent Variable Models of Inter-Regional Communication

A common approach to understanding inter-regional communication has been to identify lowdimensional latent structure that mediates the relationship between neural populations. Methods such as reduced rank regression (RRR) [23] and canonical correlation analysis (CCA) [34] identify shared and private dimensions of neural activity across regions, but leave the directionality of inter-regional communication unresolved. DLAG [22] and its multi-region extension mDLAG [26] extended this framework by modeling time-lagged latent trajectories with Gaussian process priors, enabling inference of bidirectional signal flow between neural populations. However, their reliance on stationary temporal kernels constrains them to a single, time-invariant communication mode, preventing them from capturing within-trial reconfigurations in coupling strength or direction. Dynamical system approaches, such as LFADS [19, 29] and rsLDS [18, 28] offer greater expressivity by capturing nonlinear interactions between neural populations. However, inter-regional delays must be absorbed into the latent dynamics, making directionality and lag structure difficult to disentangle from the underlying system state. Collectively, existing approaches handle either temporal delays or non-linear dynamics, but not both, leaving the problem of dynamic, directed, nonlinear inter-regional communication unresolved.

### 2.2 Transformers in Neuroscience

Transformers have been increasingly applied to neural population recordings, demonstrating advantages in generalization, scalability, and flexible mapping between neural and behavioral data [35–38]. Their application to mechanistic interpretation, however, has remained limited until recently, with transformers now being leveraged to understanding inter-regional communication. CTAE [39] introduced a transformer autoencoder that disentangles shared and private latent structure across regions and captures nonlinear and non-stationary dynamics, though it offers no insight into the directionality or temporal lags between populations. The GLM-Transformer [40] captures within-area dynamics through a transformer-based variational autoencoder, but relies on a GLM coupling term for cross-area interactions, constraining communication to be stationary. Despite these advances, the directed, dynamic nature of inter-regional communication remains unresolved, demanding new approaches.

## 3 Methods

### 3.1 Problem Formulation

Let **X** *∈* ℝ^*N ×T ×p*^ and **Y** *∈* ℝ^*N ×T ×q*^ denote simultaneously recorded spiking activity from two distinct neural populations (Fig. 1a) — population *A* with *p* units and population *B* with *q* units — measured across *N* trials of length *T* . (Note that for one application of our approach, one region is instead considered as behavioral measurements, but the above formalism still applies.) We assume that the activity of population *B* at any time *t* arises from private dynamics intrinsic to *B* and shared dynamics in the form of temporally asymmetric, bidirectional interactions with population *A* (Fig. 1b):

**Figure 1:**
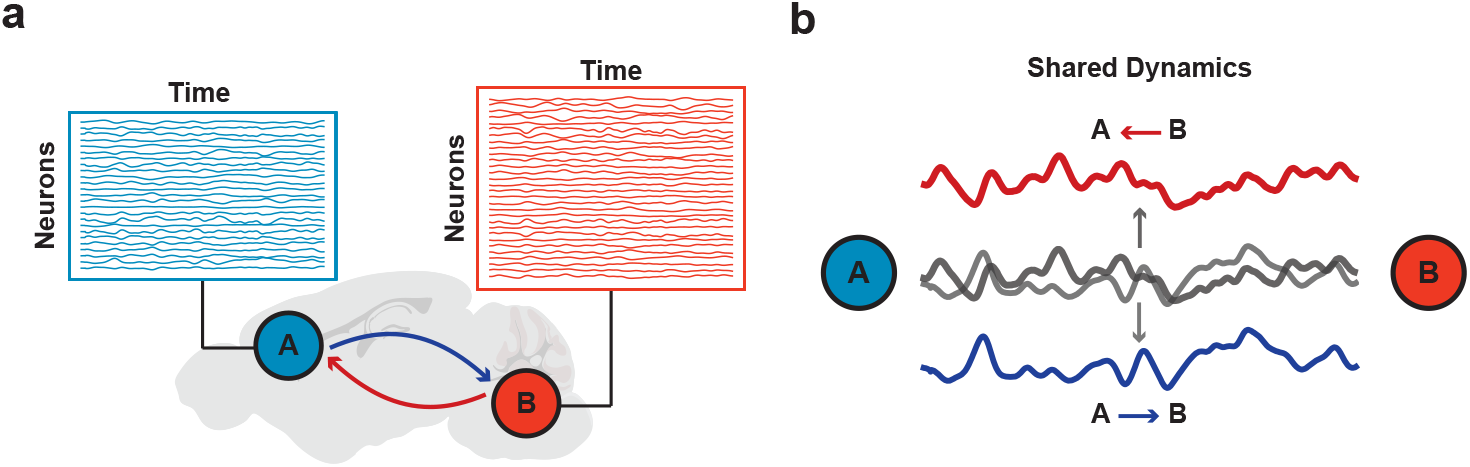
Bidirectional, concurrent interactions between neural populations. **(a)** Simultaneous recordings from two brain regions, *A* and *B*, reveal rich population activity that reflects a mixture of inter-regional signals relayed concurrently in both directions. **(b)** The shared dynamics between *A* and *B* are composed of two temporally asymmetric components: a feedforward signal (*A→ B*, blue) in which *A* leads *B*, and a feedback signal (*B→ A*, red) in which *B* leads *A*. These signals are superimposed in the recorded activity (gray), making their disentanglement a central challenge for inter-regional communication analysis.

1. **Causal (feedforward) drive**: population *A* at time *t*^*′*^ drives population *B* at a later time *t* = *t*^*′*^ + *τ*_*c*_, so that *A leads B* by delay *τ*_*c*_ *>* 0.
2. **Acausal (feedback) drive**: population *B* at time *t* is driven by population *A* at a future time *t*^*′*^ = *t* + *τ*_*a*_, so that *A lags B* by delay *τ*_*a*_ *>* 0.

The delays *τ*_*c*_ and *τ*_*a*_ are unknown, potentially distinct across communication channels, and reflect the underlying anatomical and physiological organization of the inter-regional pathways. Our goal is to jointly estimate these delays and recover the low-dimensional latent dynamics mediating each direction of communication from the simultaneously observed population recordings.

### 3.2 Model Architecture

BiXformer is a bidirectional cross-attention transformer that decomposes inter-regional communication into two parallel streams — a **causal stream** (*A→ B, A* leads) and an **acausal stream** (*B → A, B* leads) — via directionally masked cross-attention with rotary positional encoding. The full architecture is illustrated in Figure 2.

**Figure 2:**
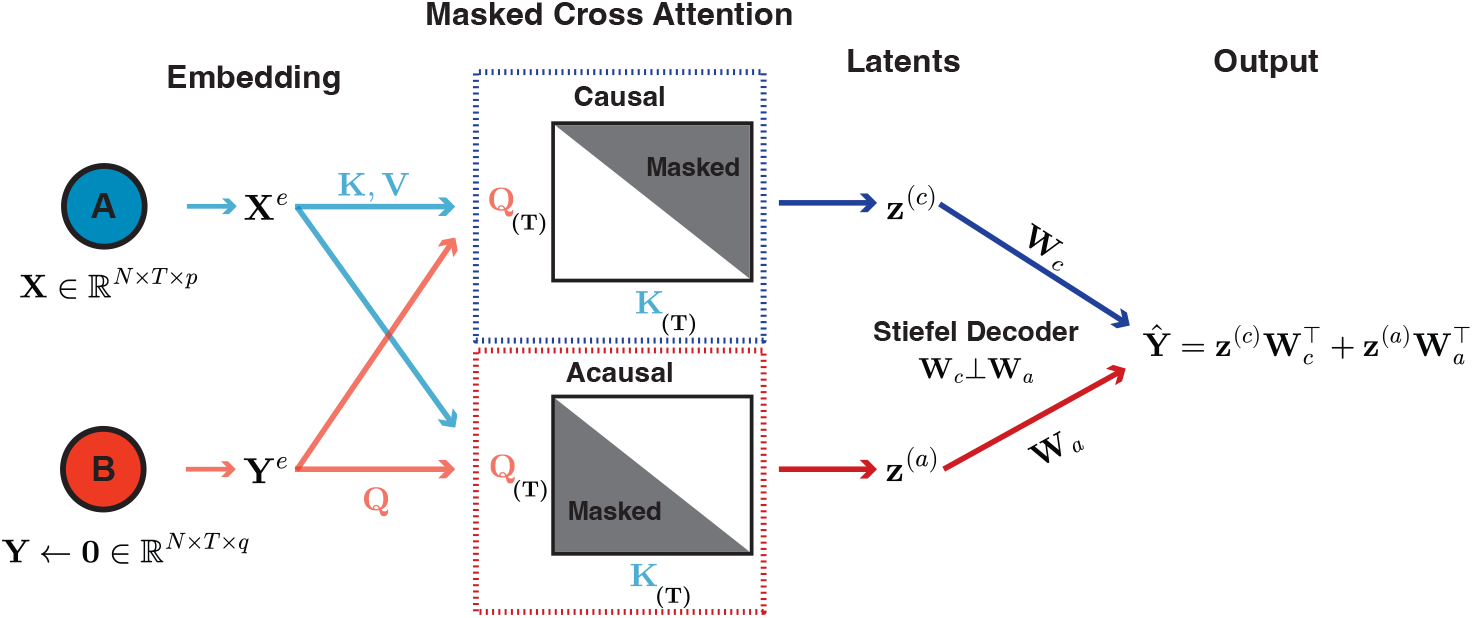
BiXformer architecture. Population *A*’s activity **X** is embedded into **X**^*e*^, providing keys and values to both streams. Population *B*’s input is zeroed during training so its embedding **Y**^*e*^ reduces to a learned bias; this bias is then linearly projected into the query matrix. Two parallel masked cross-attention modules enforce directional constraints: the causal stream attends only to past activity in *A* (*i ≥t*), and the acausal stream only to future activity (*i ≤ t*). Each stream produces a latent trajectory decoded into orthogonal subspaces of *B*’s activity space via a Stiefel-constrained decoder (**W**_*c*_ ⊥ **W**_*a*_), and their contributions are summed to reconstruct 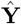.

Both populations are linearly embedded into *d*-dimensional spaces of the same dimension:

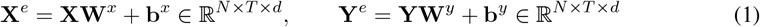

During training, the population activity of region **B** is set to zero (**Y** *←* **0**), so the embedding reduces to the learned bias alone: **Y**^*e*^ = **b**^*y*^. This bias is then projected into the query matrix **Q** = **Y**^*e*^**W**^*Q*^, so that the reconstruction of **Y** arises entirely through cross-attention to **X**. This prevents degenerate solutions in which the model bypasses the cross-attention mechanism by attending to its own input. Rotary positional embeddings (RoPE; [41]) are applied to query and key vectors so that attention scores depend only on the relative temporal offset *t− t*^*′*^, enabling each head to learn a preference for a specific inter-regional lag.

#### 3.2.1 Masked Cross-Attention Streams

BiXformer uses two parallel cross-attention modules, each with *H/*2 heads (*H* total). In both streams, queries are formed from population *B*’s embedding, while keys and values are formed from population *A*’s embedding:

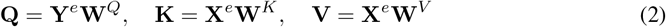

The masked attention weight from population *B* at time *t* to population *A* at time *i* is:

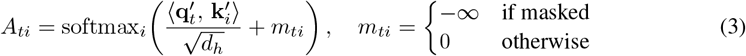

where setting *m*_*ti*_ = *−∞* drives the corresponding attention weight to exactly zero after the softmax, effectively removing that position from the attended context. This is used to enforce directional constraints in each stream:

##### Causal stream

positions *i > t* are masked, so population *B* at time *t* attends only to population *A* at times *i ≤ t*, enforcing that *B*’s activity is explained only by past or concurrent activity in *A*, i.e., *A* leads *B*.

##### Acausal stream

positions *i < t* are masked, so population *B* at time *t* attends only to population *A* at times *i ≥ t*, enforcing that *B*’s activity is explained only by future or concurrent activity in *A*, i.e., *B* leads *A*.

The inter-regional communication delay is read out from the **diagonal profile** of each head’s *T × T* attention matrix — the mean attention weight as a function of lag *δ* = *i − t*:

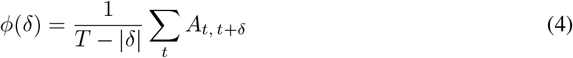

The profile concentrates at *δ*^*\**^ = arg max_*δ*_ *ϕ*(*δ*), providing a direct, interpretable estimate of the lag preferred by that communication channel. After multi-head aggregation and a linear out-projection, each stream output is compressed to an *r*-dimensional latent representation:

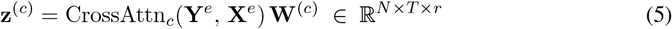

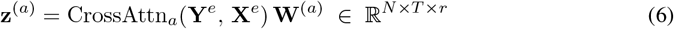

#### 3.2.2 Stream-Orthogonal Stiefel Decoder

To guarantee that causal and acausal streams decode into non-overlapping subspaces of population *B*’s activity space, we impose a **hard orthogonality constraint** on the decoder weight matrices via QR reparameterization [42]. Given learnable raw parameters 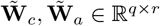, we compute the thin QR decomposition of their column-concatenation at every forward pass:

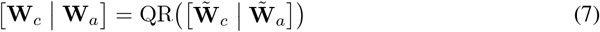

The resulting matrices **W**_*c*_, **W**_*a*_ ∈ ℝ^*q×r*^ satisfy 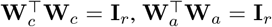, and 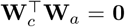, constraining the two decoders to lie on the Stiefel manifold and to span orthogonal subspaces of ℝ^*q*^ . Population *B*’s activity is reconstructed as the sum of the two stream contributions:

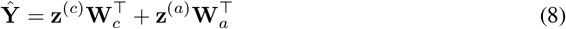

This hard constraint guarantees exact subspace separation at every forward pass, without requiring tuning of an orthogonality penalty weight.

### 3.3 Training Objective

BiXformer is trained to minimize the mean-squared reconstruction error of target population *B*:

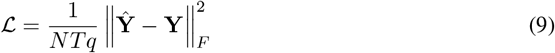

The Stiefel orthogonality constraint on the decoder (**W**_*c*_ ⊥ **W**_*a*_) is enforced structurally, ensuring that the causal and acausal streams write into orthogonal subspaces of the target population’s activity. This means that the contribution of each stream to any given target variable can be read off directly and unambiguously, without mixing. Directional separation between streams is enforced by the causal masks, so no auxiliary losses are required. Parameters are optimized with Adam, with an 80–20 train-test split; training epochs vary over experiment; model performance is reported on the held-out fold of each split (see Appendix A1).

## 4 Experiments

### 4.1 Validating BiXformer on Synthetic Data

To verify that BiXformer correctly recovers inter-regional delays and latent communication structure, we generated synthetic datasets with known ground truth using Gaussian Process (GP)-driven latent dynamics (Fig. 3a; see Appendix B1). The dataset was designed to reflect the scale and structure of real neural recordings: two populations of *p* = 80 and *q* = 50 neurons recorded over 200 trials of *T* = 200 time bins (5 ms resolution), with both shared cross-area latent dimensions and population-specific private latents to model within-area dynamics not involved in inter-regional communication. Four cross-area latent dimensions were included — two causal (*A →B*) and two acausal (*B →A*) — each with GP timescales and communication delays independently sampled from a uniform distribution spanning a range of biologically plausible values. Population activity was constructed by multiplying the latent trajectories by random mixing matrices and adding i.i.d. Gaussian observation noise. To ensure clean separation between shared and private variance in the ground truth, the within-area loading vectors were projected onto the null space of the cross-area loading matrix (via SVD), guaranteeing that private latents explain variance exclusively in directions orthogonal to those used by cross-area communication.

**Figure 3:**
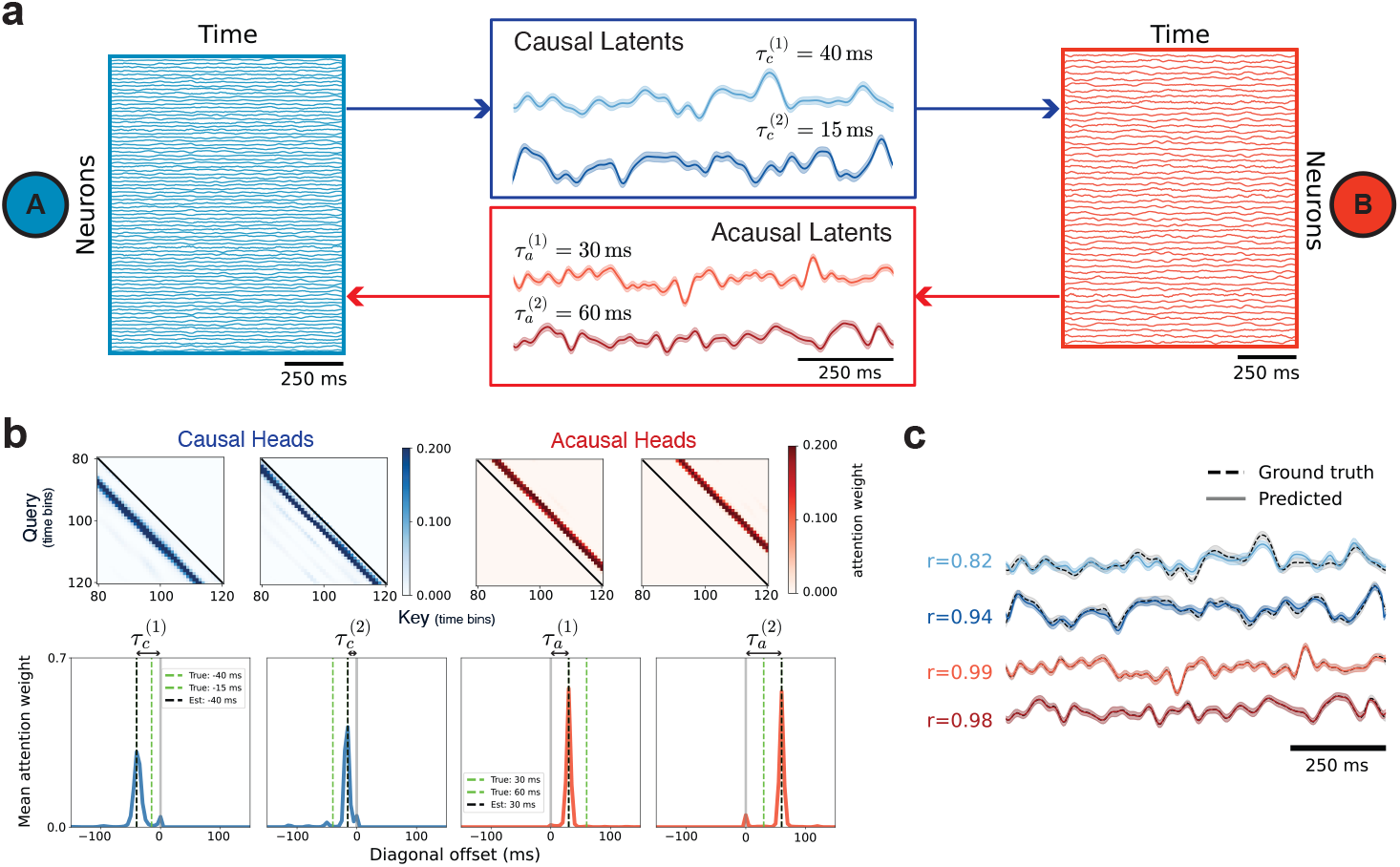
BiXformer recovers directed latent dynamics and communication delays from synthetic data. **(a)** Population activity from regions *A* (blue) and *B* (red) is decomposed into causal (*A →B*) and acausal (*B →A*) latents, each associated with a known ground-truth delay. **(b–c)** Results are shown for a representative held-out fold from 5-fold cross-validation. **(b)** Representative attention heads with the strongest diagonal profiles are shown (zoomed in for clarity); diagonal profiles peak at offsets closely matching the true delays (dashed lines). See Appendix D1 for all heads. **(c)** Predicted latent trajectories (solid) closely track ground-truth latents (dashed) across all four dimensions (Pearson *r* reported for single-trial predictions).

Following training, we evaluated BiXformer on held-out data. A subset of attention heads developed peaked diagonal profiles at offsets closely matching the true communication delays (Fig. 3b), while the remaining heads exhibited diffuse, non-selective attention patterns — consistent with the model allocating representational capacity selectively across channels (see Appendix D1). Latent trajectories recovered by each stream were well-correlated with the corresponding ground-truth GP signals on single trials (Fig. 3c).

To compare our approach to established methods, we evaluated BiXformer on synthetic data generated by DLAG’s generative model, demonstrating that BiXformer accurately recovers inter-regional communication delays under the same ground-truth conditions used to validate DLAG, while achieving higher predictive performance and substantially faster training (Appendix C). Together, these results establish that BiXformer reliably identifies the direction, magnitude, and latent structure of bidirectional inter-regional communication in synthetic data.

### 4.2 Disentangling Feedforward and Feedback Signals in Neural-Behavioral Recordings

As described in the Introduction, neural activity in sensorimotor regions simultaneously reflects feedforward motor commands and ascending sensory feedback, which are temporally offset yet super-imposed within the same population — creating a confound that motivates an explicitly bidirectional decomposition.

We applied BiXformer to simultaneous electrophysiological recordings (118 units; spike counts binned at 2.5 ms) from the IRN/PARN of the medulla and high-speed videography (56 orofacial kinematic keypoint trajectories) in a head-fixed mouse performing a delayed licking task (Fig. 4a). Although our approach is designed primarily for inter-regional communication (as we do below), application to a neural-behavioral dataset enables greater interpretability owing to the direct interpretation that behavior affords. The medulla exhibits highly stereotyped, rhythmic activity tightly coupled to orofacial movement, reflecting its prominent role in motor execution — making it a compelling model system for evaluating bidirectional decomposition (Fig. 4b). To aid interpretability and visualization, PCA was applied independently to both behavioral and neural datasets; all principal components were retained, preserving 100% of the variance.

**Figure 4:**
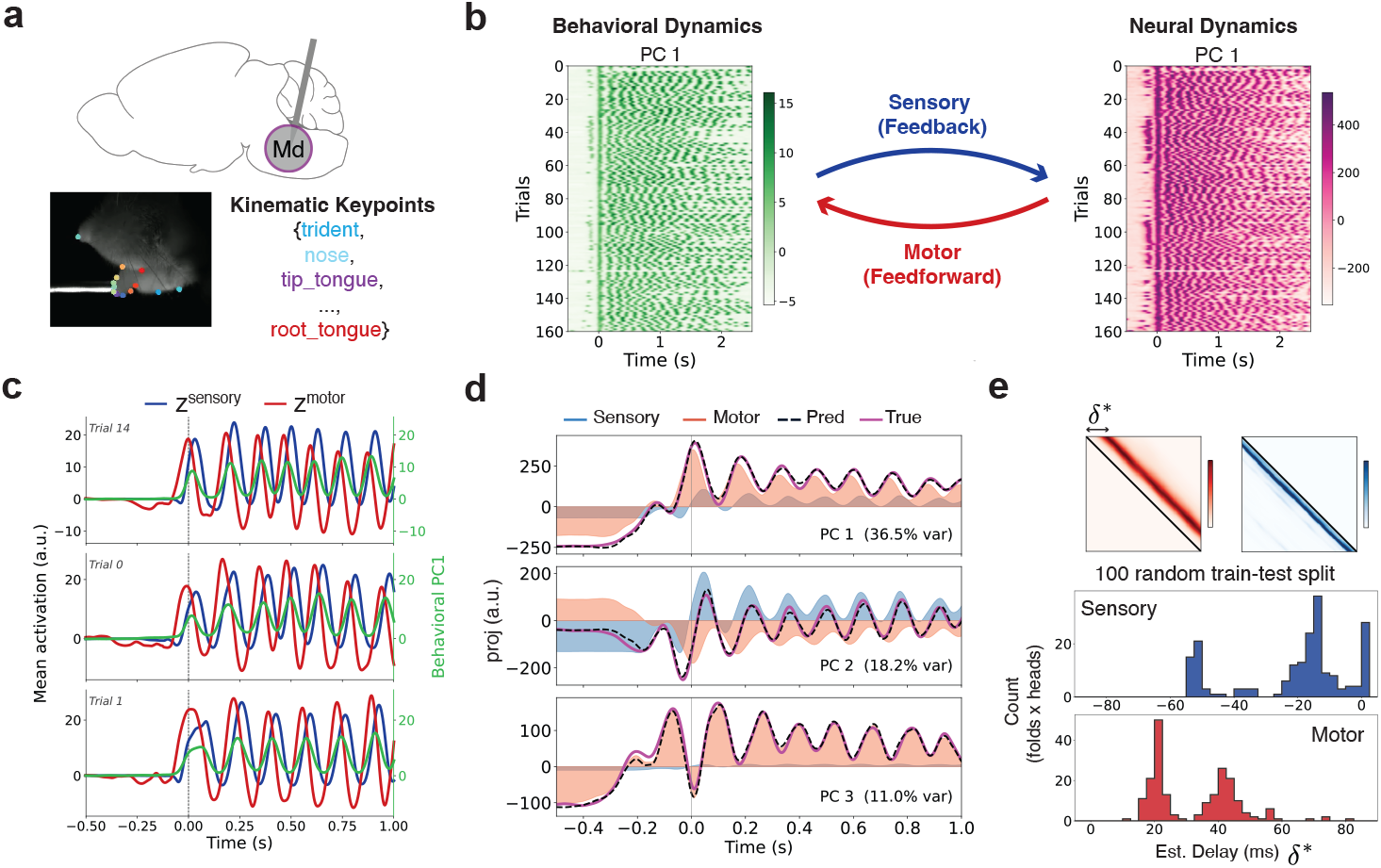
BiXformer disentangles sensory and motor signals in neural-behavioral recordings from the medulla. **(a)** Electrophysiological recordings from the medulla (Md) (sagittal view) during a rhythmic licking task, with orofacial kinematics tracked via high-speed videography. **(b)** The first principal component of behavioral keypoint features (left) and Md population activity (right) for multiple behavioral trials; time is aligned to first lickport contact. Bidirectional arrows indicate the sensory (feedback, blue) and motor (feedforward, red) pathways modeled by BiXformer. **(c)** Mean activation of sensory (*z*^sensory^) and motor (*z*^motor^) latents, overlaid with behavioral PC1 (40.4% variance explained) for multiple trials. **(d)** Contribution of each stream to reconstructing the top neural PCs (mean). **(e)** Estimated communication delays across 100 random train-test splits (2 heads per stream per run).

Evaluated on held-out data, BiXformer successfully disentangles feedforward and feedback signals within medulla dynamics (predicted using behavioral dynamics). The sensory (*z*^sensory^) and motor (*z*^motor^) latents exhibit distinct, interpretable temporal profiles: the motor stream precedes the dominant behavioral PC (40.4% of variance explained) consistent with a feedforward motor command, while the sensory stream follows it, consistent with ascending sensory feedback (Fig. 4c). Despite this explicit disentanglement, the model retains strong predictive power, achieving *R*^2^ = 0.645*±* 0.023 across 5-fold cross-validation (see Appendix D2).

The Stiefel orthogonality constraint on the decoder (Fig. 2) ensures that the two streams write into orthogonal subspaces of the medulla’s neural population activity, allowing us to directly attribute each neural PC’s variance to either the sensory or motor stream (Fig. 4d). We find that the motor stream strongly dominates the first PC of medulla activity, consistent with the IRN/PARN being a major motor hub [43, 44]. Finally, to characterize the temporal structure of communication within each stream, we estimated neural-behavioral lags from the attention heads across 100 random train-test splits (2 heads per stream per run; Appendix D3). The resulting delay distributions reveal distinct characteristic lags within each stream (Fig. 4e), suggesting that multiple signals are transmitted at different timescales within both the sensory and motor pathways (GMM fit: sensory *µ* = {16.0, 50.5 } ms, *σ* = {5.2, 6.2} ms; motor *µ* = {19.4, 40.4} ms, *σ* = {2.6, 3.6} ms; see Appendix D4). Across sessions from two different animals, the estimated delay distributions remain distinct but shifted in their characteristic lags (see Appendix D5), reflecting differences in recording sites and the specific neural populations sampled within the IRN/PARN. Together, these results demonstrate that BiXformer can recover interpretable feedforward and feedback components from mixed neural-behavioral recordings, revealing the temporal structure of communication within each pathway, while preserving strong predictive performance.

### 4.3 Uncovering Directed Interactions Between S1 and M1 During a Delayed Reward Task

Next, we used BiXformer to disentangle sensory and motor related signals from multi-site neural recordings during movement. Recent experimental measurements from primary motor and sensory areas from mice executing controlled movements has demonstrated a rich interplay of movement and sensory related signals across the cortex [15]. Activity in primary motor cortex (M1) was shown to contain temporally multiplexed sensory and motor related signals, each of which was differentially represented during specific temporal epochs during a licking task. Motor signals were transiently present during periods when the consequences of movements were uncertain while “residual” low-amplitude sensory signals were present during all periods of the task. Primary somatosensory cortex (S1) was active throughout the movement period, suggesting that residual M1 activity may reflect ongoing sensory input from S1 rather than intrinsic motor dynamics.

To test this hypothesis, we applied BiXformer to simultaneous S1 and M1 recordings, treating S1 as the source population (*X*) from which to predict M1 activity (*Y*), thereby decomposing M1 activity into components attributable to S1 input (S1 *→* M1) and components that precede S1 activity and are thus not attributable to it (M1 *→* S1) (Fig. 5a).

**Figure 5:**
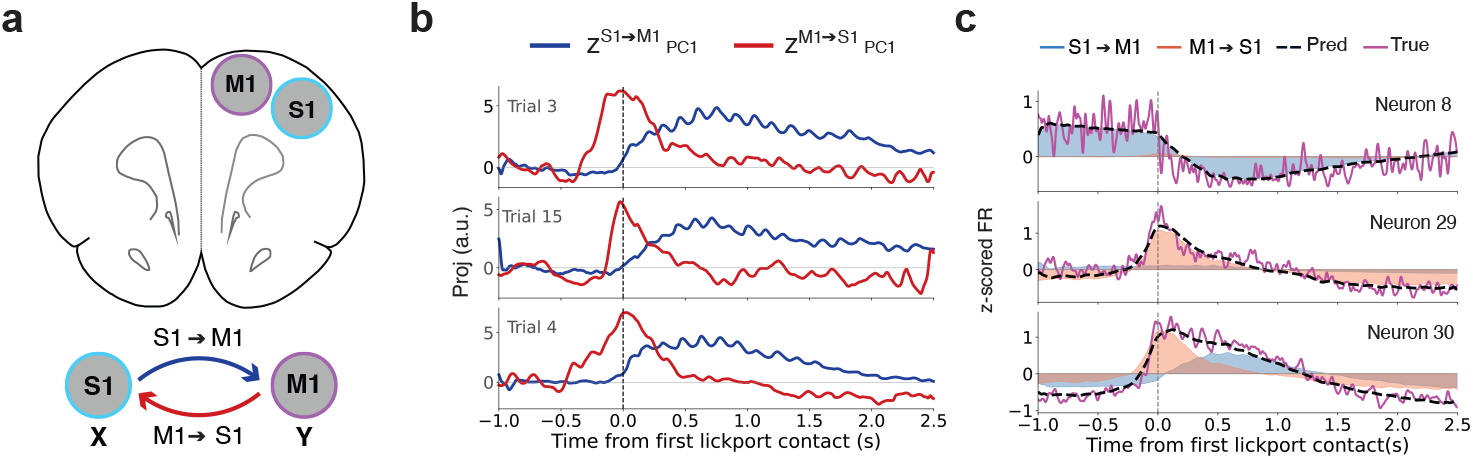
BiXformer disentangles bi-directional signals in simultaneous S1–M1 recordings during a delayed reward task. **(a)** BiXformer is applied to simultaneous recordings in S1 (*X*) and M1 (*Y*), where the causal stream models S1 *→*M1 communication and the acausal stream models M1 *→*S1 communication. **(b)** PC1 of the S1 *→*M1 (*z*^S1*→*M1^) and M1 *→*S1 (*z*^M1*→*S1^) latent spaces, capturing 51.0% and 89.9% of their respective variances, shown for three example trials. **(c)** Contribution of each stream to reconstructing the mean activity of individual neurons; three example neurons shown.

The latent dynamics recovered by each stream show distinct signatures of S1-M1 communication. The dominant mode of the S1 *→*M1 stream (PC1, capturing 51.0% of variance) exhibits a sustained signal that rises around the first lickport contact and persists throughout the lick bout, consistent with an ongoing sensory signal transmitted from S1 to M1 (Fig. 5b). The dominant mode of the M1 *→*S1 stream (PC1, capturing 89.9% of variance) resembles the identified engagement-disengagement profile that peaks sharply at first lickport contact and rapidly dissipates [15]. These results demonstrate the ability of BiXformer to partition neural responses into motor- and sensory-related signals originating within interconnected neural populations.

Assessing each stream’s contribution to reconstructing individual M1 neuron activity revealed heterogeneity across the population, reflecting the diversity of functional signals present in M1 during movement. (Fig. 5c). Some neurons were well-described by the M1*→* S1 stream alone. For example, Neuron 29 exclusively exhibited a strong motor drive that disengaged rapidly, and reflected virtually no sensory-related signals from S1. Other neurons show mixed contributions from both streams (e.g, Neuron 30), consistent with the highly mixed responses observed across sensorimotor circuits and which are necessary for sensory-guided motor control. Somewhat surprisingly, a subset of neurons were predominantly and almost exclusively reconstructed by the S1 *→*M1 stream (e.g, Neuron 8), indicating they were largely and almost exclusively driven by S1 input. These neurons did not reflect strong motor related signals around movement initiation, suggesting a different functional contribution to motor control than neurons primarily reflecting M1 *→*S1 signals. Surprisingly, these neurons were active prior to movement onset, suggesting the S1*→* M1 pathway may carry more than purely sensory-driven signals, such as anticipatory or top-down influences during motor planning. Collectively, these findings are consistent with the hypothesis that S1 input contributes to residual M1 activity following cortical disengagement, while also revealing intriguing top-down sensory signals that provide deeper insight into the nature of S1-M1 communication.

## 5 Discussion

We introduced BiXformer, a transformer that disentangles directed, time-delayed communication from simultaneous neural population recordings. Masked cross-attention enforces directionality and a Stiefel-constrained decoder enforces subspace separation, yielding latent dynamics that are both expressive and structurally interpretable, without linearity or stationarity assumptions. On synthetic data, BiXformer recovers latent trajectories and inter-regional delays. Applied to medulla recordings during rhythmic licking, it cleanly separates feedforward motor commands from ascending sensory feedback. Applied to simultaneous S1–M1 recordings during a delayed reward task, it uncovered a heterogeneous M1 population differentially driven by S1 input versus intrinsic motor dynamics, including a subset whose pre-movement activity is explained by the S1*→*M1 stream.

Existing approaches to inter-regional communication face a recurring tradeoff between expressivity and interpretability. Linear-Gaussian methods [22, 26] model delays explicitly through time-shifted latent Gaussian processes, but their linear observation model precludes nonlinear interactions between populations. Dynamical-system approaches [19, 18] capture nonlinear dynamics but represent delays only implicitly through the latent state, obscuring the direction and timing of communication. Recent transformer-based methods either omit directionality [39] or constrain communication to be stationary [40]. BiXformer resolves this tradeoff: masked cross-attention enforces directionality architecturally, per-head attention expose lag structure interpretably, and the underlying mapping remains nonlinear.

### Limitations & Future Directions

Our experiments involve only two populations at a time. Extending BiXformer to a true multi-region setting — with directed communication estimated across many populations — is conceptually straightforward via additional cross-attention streams, but raises non-trivial questions about identifiability, scaling, and interpretation of inter-regional interactions. We currently fix the number of attention heads per stream and observe empirically that only a subset develop selective diagonal profiles. A principled head-pruning procedure — or a sparsity-inducing prior over heads — would yield more parsimonious models and sharper estimates of the number of distinct communication channels. Finally, the latent dimensions are unconstrained beyond their decoder geometry; imposing temporal or population-level sparsity would likely improve identifiability of individual communication channels and aid interpretation, particularly when distinct channels operate at different timescales. Finally, the diagonal profile of each attention head provides a natural primitive for further architectural engineering: heads could be parameterized to learn unimodal lag kernels with explicit centers and widths, yielding a more compact and directly interpretable representation of communication delays.

## Acknowledgments

This work was supported by NIH/NINDS R01NS121409 (MNE), NIH/BRAIN U19NS137920 (MNE), NSF CAREER 2239412 (MNE), and a Scialog grant from the Research Corporation for Science Advancement (BD).

## Appendix

### A Model Details

#### A.1 Training Details

All models were trained using the Adam optimizer [45] with gradient norm clipping at 0.5. For the synthetic data experiments, BiXformer was trained for 500 epochs per fold under 5-fold cross-validation. For the neural-behavioural (IRN) real-data experiments, training ran for 400 epochs per fold, evaluating 5 of 10 folds from a 10-fold cross-validation split. For the multi-region (S1–M1) experiments, training ran for 400 epochs per fold under standard 5-fold cross-validation. For the 100-run random train–test split evaluation used to assess delay distribution robustness (Figures 4e and D5), each independent run was trained for 400 epochs on an 80%*/*20% random train–test split of the data, with the split seed set equal to the run index for reproducibility.

#### A.2 Hyperparameters

No hyperparameter search was conducted; all values were set based on initial exploratory runs and held fixed across experiments. Table 1 reports the hyperparameters used across all experiments. The synthetic and real-data experiments share the same model dimension, batch size, and learning rate, with minor differences in the number of attention heads and latent dimensionality.

**Table 1:**
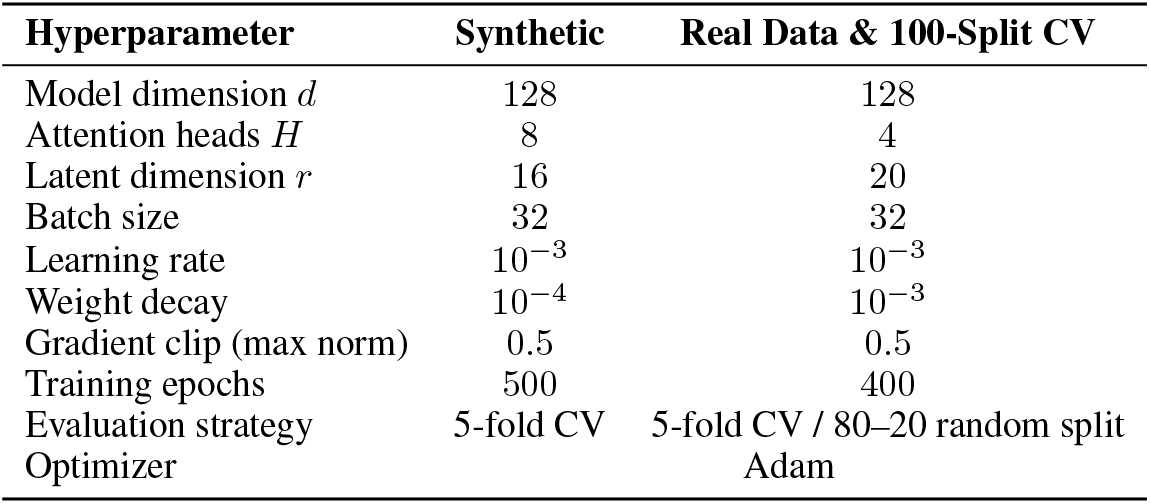
Hyperparameters used across all experiments.

#### A.3 Compute

Synthetic and real-data cross-validation experiments were run on an Apple Silicon GPU (via MPS). The 100-run random train–test split evaluation was conducted across three sessions from three animals, distributed across NVIDIA A100, A40, and L40S GPUs.

### B Datasets

#### B.1 Synthetic Data Generation

We generated synthetic paired population recordings **X**^(*n*)^ *∈* ℝ^*T ×p*^ and **Y**^(*n*)^ *∈* ℝ^*T ×q*^ for *n* = 1, *…, N* trials, where *T* is the number of time bins and *p, q* are the dimensionalities of the two populations.

##### Latent processes

Each latent *z*_*k*_ is an independent draw from a Gaussian Process (GP) with a squared-exponential covariance kernel,

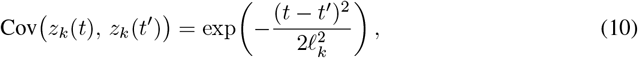

where *l*_*k*_ *>* 0 is the GP timescale of latent *k*, controlling the smoothness of the trajectory over time.

##### Cross-area latents and delays

There are *n*_*c*_ causal latents (**X** leads **Y**, delay *τ*_*k*_ *>* 0) and *n*_*a*_ acausal latents (**Y** leads **X**, delay *τ*_*k*_ *<* 0), where *τ*_*k*_ denotes the inter-regional communication delay for latent *k*.

##### Private latents

In addition, *n*_*x*_ within-**X** private latents 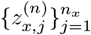 and *n*_*y*_ within-**Y** private latents 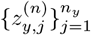 are drawn from independent GPs and contribute to only one population with no delay.

##### Observations

Observations are formed by linearly mixing the latents and adding i.i.d. Gaussian observation noise ***ε*** *∼ 𝒩* (**0**, *σ*^2^**I**):

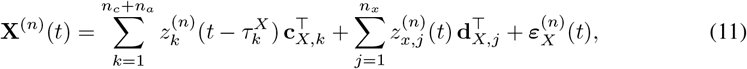

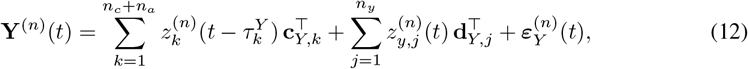

where 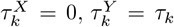 for causal latents (so **Y** is shifted by delay *τ*_*k*_ *>* 0 relative to **X**) and 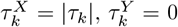 for acausal latents (so **X** is shifted relative to **Y**). The vectors **c**_*X,k*_ *∈* ℝ^*p*^ and **c**_*Y,k*_ *∈* ℝ^*q*^ are fixed random cross-area loading vectors. The private loading vectors **d**_*X,j*_ *∈* ℝ^*p*^ and **d**_*Y,j*_ *∈* ℝ^*q*^ are projected onto the null space of the cross-area loading matrix, ensuring that private variance is orthogonal to the shared cross-area subspace.

##### Demo dataset

The dataset used in Section 4.1 was generated with *N* = 200 trials, *T* = 200 bins (Δ*t* = 5 ms), *p* = 80, *q* = 50, *n*_*c*_ = *n*_*a*_ = 2, *n*_*x*_ = *n*_*y*_ = 2. True communication delays were ***τ*** = [+40, +15, *−* 30, *−* 60] ms (positive = **X** leads **Y**; negative = **Y** leads **X**) and cross-area GP timescales were ***l***_cross_ = [25, 20, 15, 18] ms (causal: *l* = 25, 20 ms; acausal: *l* = 15, 18 ms), private-**X** timescales ***l***_*x*_ = [25, 12] ms, and private-**Y** timescales ***l***_*y*_ = [30, 18] ms.

#### B.2 Neural-Behavioral Recordings (IRN)

Adult C57BL/6J mice were implanted with a custom metal headbar, water-restricted, and trained on a head-fixed licking task in which the animal produced tongue movements across a range of angles toward a lickport to obtain water reward, with both go-cue and self-initiated trial types interleaved within each session. After training, a small craniotomy was made over the medullary reticular formation, and acute extracellular recordings were performed using 4-shank Neuropixels 2.0 probes (IMEC) targeting the intermediate reticular nucleus and adjacent areas in which putative licking oscillator neurons have been described [43, 44]. Voltage signals were acquired at 30 kHz using SpikeGLX, band-pass filtered between 300 Hz and 6 kHz, spike-sorted with Kilosort 2.5, and automatically curated based on standard quality metrics. Spike times were binned at 2.5 ms resolution and the resulting firing rates were smoothed with a Gaussian kernel (*σ* = 20 ms) prior to analysis. During recording, high-speed video of the orofacial region was acquired at 400 Hz using bottom-view and side-view cameras, and tongue and jaw keypoints were tracked with DeepLabCut.

Three recording sessions from three animals were analysed, yielding 118, 162, and 181 isolated units respectively. Orofacial kinematics were tracked at 56 keypoints across all sessions.

#### B.3 Multi-Region Neural Recordings (S1–M1)

Data were obtained from [15]; full surgical and experimental details are provided therein. Briefly, adult C57BL/6J mice (8–13 weeks) were implanted with a headbar and trained on a Delayed Reward licking task. A craniotomy was made over the tongue region of primary motor cortex (tjM1; AP: +2.2 mm, ML: *±*2.5 mm) and tongue-jaw primary somatosensory cortex (tjS1; AP: +0.5 mm, ML: *±*3.5 mm), and simultaneous extracellular recordings were performed in both areas using Neuropixels 1.0 probes. Spike times were binned at 2.5 ms resolution and firing rates smoothed with a Gaussian kernel (*σ* = 10 ms), yielding 47 units in tjM1 and 18 units in tjS1.

In the Delayed Reward task, each trial began with a 2 s LED cue followed by an auditory Go Cue, after which the animal licked a motorized port to receive a water reward. On each trial, reward was delivered on either the first (R1) or fourth (R4) port contact with equal probability (50%), with the lickport cycling among three lateral positions in blocks of 10 trials. Only R4 trials were used in the analysis presented here.

### C Model Comparison

#### C.1 DLAG’s Generated Dataset

We generated synthetic data using DLAG’s own simulation framework [22], via the script `dlag_groundtruth_data.m` from their public repository. The dataset consists of 200 trials of *T* = 100 time bins (Δ*t* = 1 ms), with 10 observed neurons in each of two areas and 4 across-area latents with ground-truth GP timescales ***l*** *≈* [7.0, 9.5, 6.9, 6.7] ms and delays ***τ*** *≈* [+12.7, +1.7, *−*5.5, *−*12.0] ms (positive = area 1 leads area 2).

#### C.2 Delay Estimation and Predictive Performance

We fit DLAG to this dataset across 5 independent random seeds and fit BiXformer via 5-fold cross-validation on the same data. Delay estimation accuracy was quantified by matching estimated delays to ground-truth values via the Hungarian algorithm (per fold) and reporting mean absolute error (MAE, in ms). Predictive performance was measured as variance-weighted *R*^2^ on held-out trials. BiXformer achieved lower delay MAE (2.8 ms vs. 5.8 ms) and higher held-out *R*^2^ (0.332 *±* 0.026 vs. 0.273 *±* 0.019).

**Figure 6:**
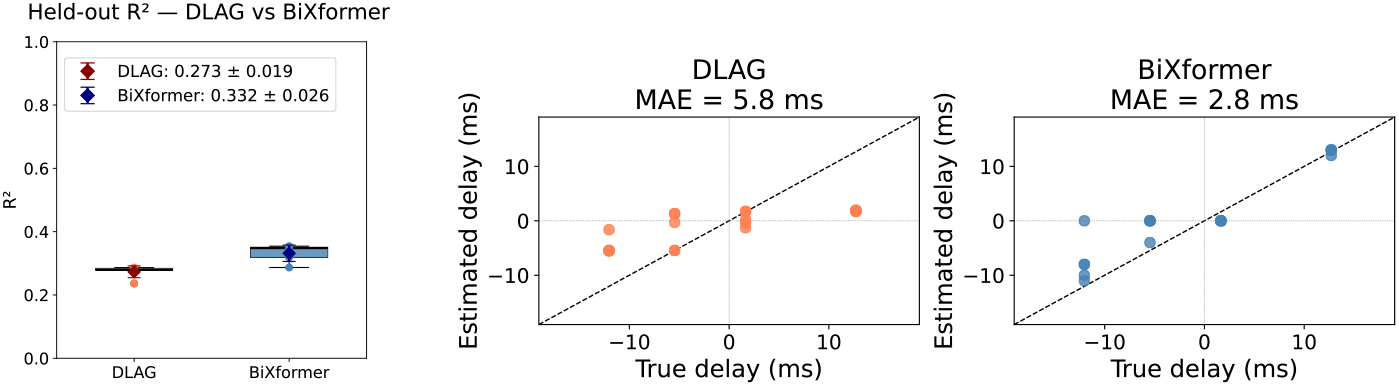
Comparison of BiXformer and DLAG on synthetic data. *Left:* Held-out *R*^2^ across 5 independent DLAG runs and 5 BiXformer cross-validation folds (box: interquartile range; whiskers: full range; diamond: mean*±* std). BiXformer explains more variance than DLAG (0.332 *±*0.026 vs. 0.273 *±* 0.019). *Center and Right:* Estimated vs. ground-truth inter-area delays for DLAG and BiXformer, respectively. Each point is one delay matched via the Hungarian algorithm. BiXformer achieves lower mean absolute error (MAE = 2.8 ms vs. 5.8 ms), recovering both the sign and magnitude of all four delays more accurately.

#### C.3 Computational Cost

On this dataset, DLAG required on average 21.7 minutes per run (5 seeds; trained until convergence or up to 5,000 EM iterations), while BiXformer completed each fold in approximately 1.2 minutes (5 folds; 1,000 epochs each), a roughly 18*×* reduction in wall-clock time. We note that these runtimes are not directly comparable: DLAG was run on CPU, while BiXformer was run on an Apple Silicon GPU (via MPS). Nonetheless, the difference reflects the practical training overhead of each approach on the hardware it was designed to use.

### D Additional Results

#### D.1 Synthetic Data: All Attention Heads, Diagonal Profiles, and Performance Across Folds

**Figure 7:**
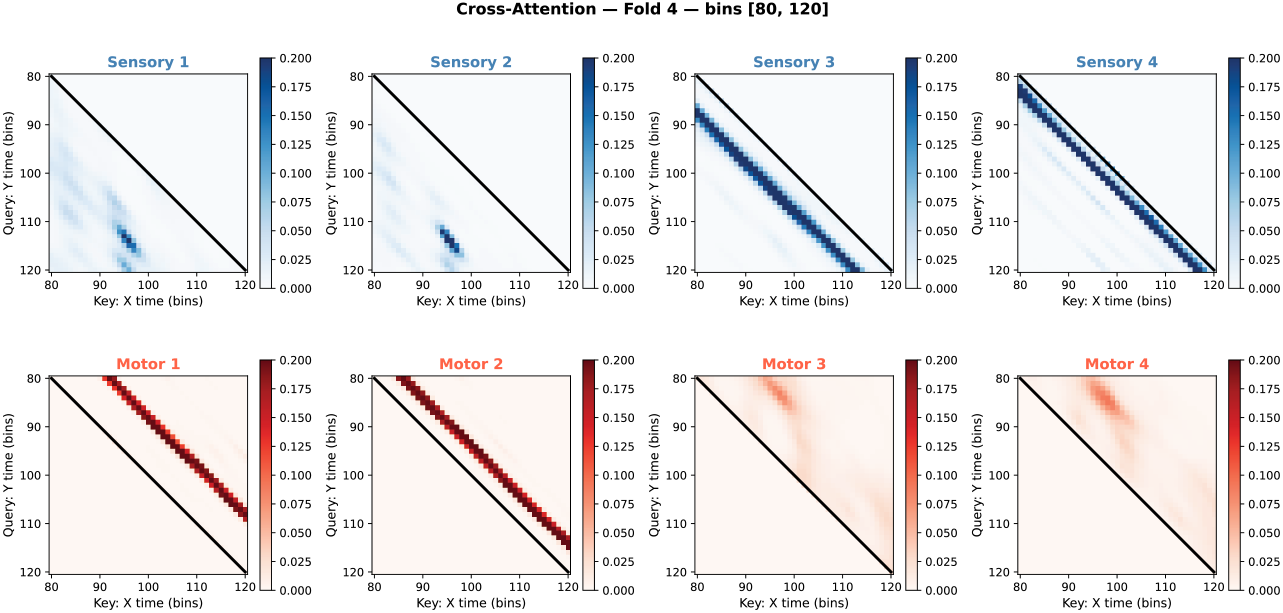
Cross-attention weight maps. Trial averaged across all eight heads — four causal (blue) and four acausal (red) — for an example fold of the synthetic experiment, shown over a zoomed time window. Two heads of each stream display sharp off-diagonal bands consistent with the ground-truth delays; the remaining heads are diffuse.

**Figure 8:**
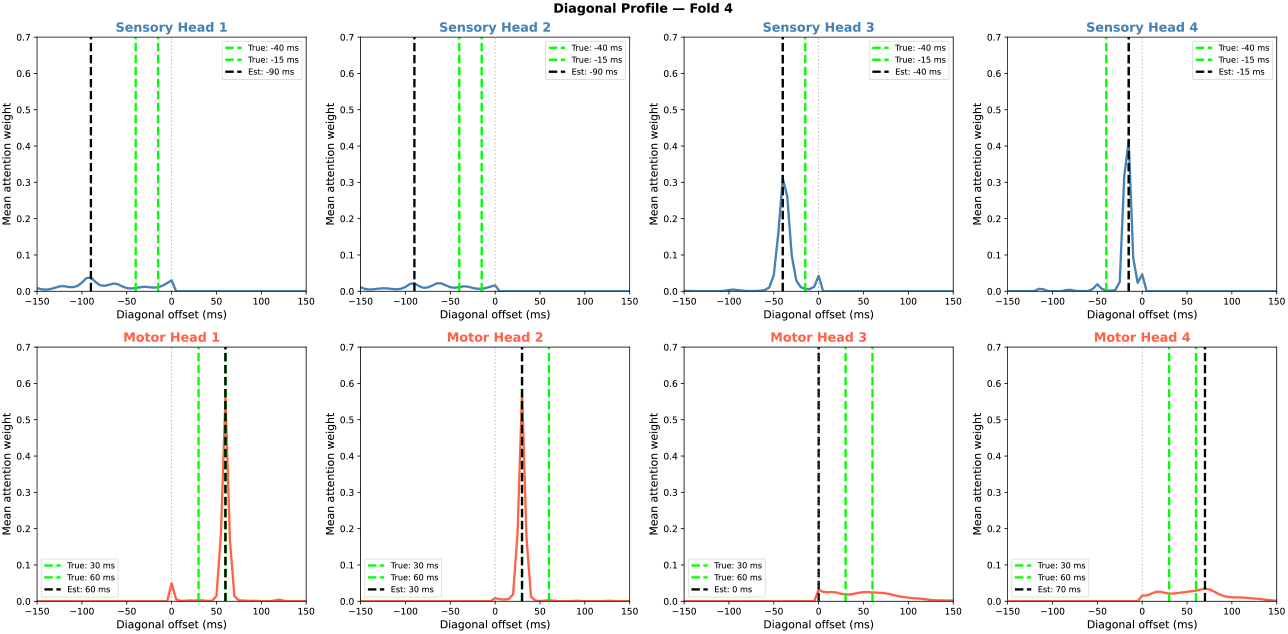
Diagonal profiles. Trial averaged across all eight heads. Causal heads 3 and 4 accurately recover the ground-truth delays of *−*40 and *−*15, ms, respectively; causal heads 1–2 yield broad, uninformative profiles. Acausal heads 1 and 2 recover +60 and +30, ms; heads 3–4 are diffuse. Green dashed lines mark ground-truth delays; black dashed lines indicate the per-head peak estimate.

#### D.2 Constrained vs. Unconstrained Model: Reconstruction Performance Across Five Folds

**Figure 9:**
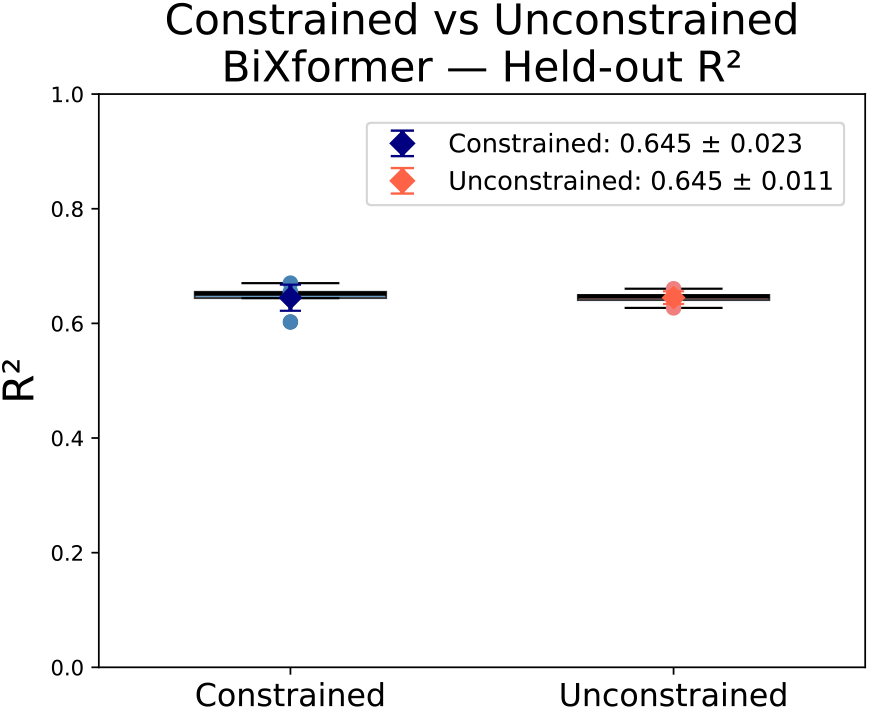
Held-out *R*^2^ for the constrained BiXformer and an unconstrained variant across 5 cross-validation folds on the neural-behavioural (IRN) dataset. The unconstrained model removes both the causal/acausal attention masking and the Stiefel manifold constraint on the decoder, while keeping all other hyperparameters and parameter counts identical. Boxes show the interquartile range; individual fold values are overlaid as points; diamonds with error bars indicate mean *±* standard deviation.

#### D.3 Attention Heads and Diagonal Profiles for Neural-Behavioral Experiment

**Figure 10:**
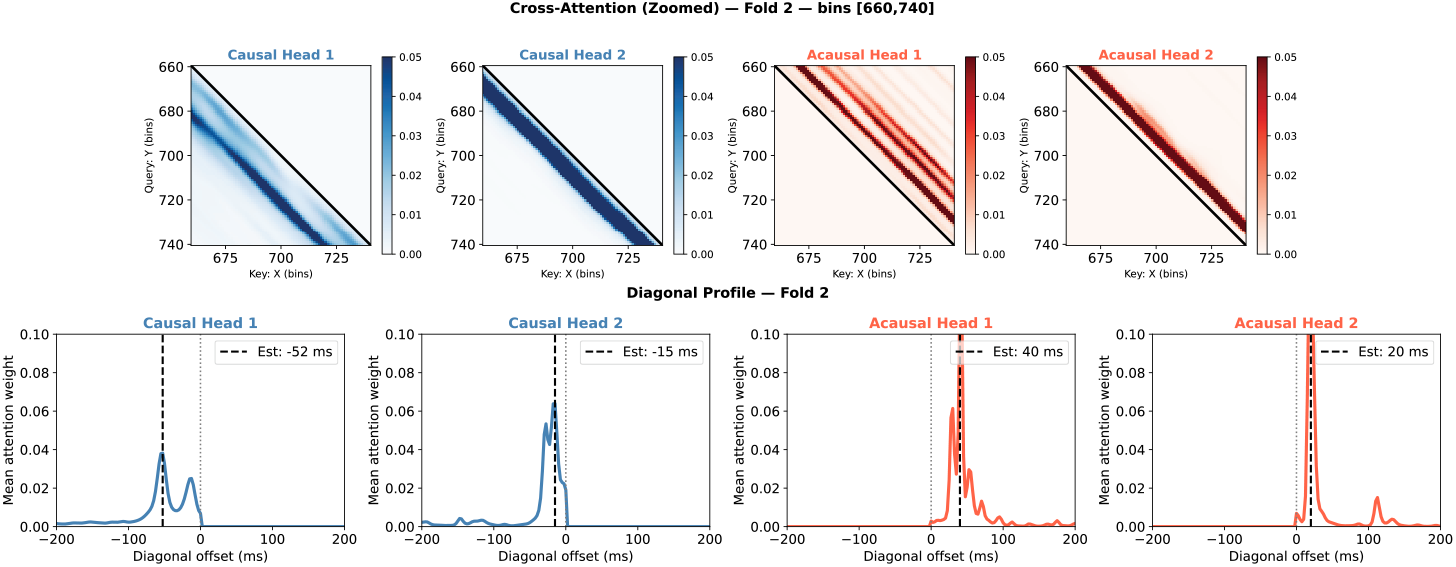
Attention maps (top) and diagonal profiles (bottom) for an example fold of the neural-behavioral experiment. Causal heads identify negative diagonal offsets (*−*52 and *−*15 ms), indicating that behavior leads neural activity, while acausal heads identify positive offsets (+40 and +20 ms), indicating that neural activity leads behavior. The sharp, well-localized peaks in all four profiles demonstrate that the model consistently latches onto specific temporal lags between medullary neural activity and orofacial kinematics.

#### D.4 GMM Model Fit

**Figure 11:**
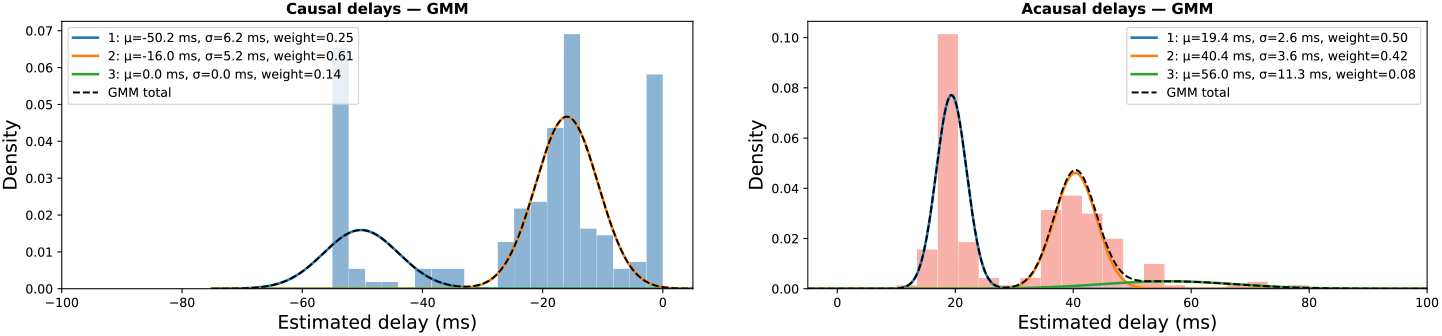
Gaussian Mixture Model (GMM) fit to the distribution of estimated delays across 100 random 80/20 train–test splits for causal (top) and acausal (bottom) heads. For each head, delays were pooled across all splits and fit using `sklearn.mixture.GaussianMixture` with full covariance. The number of components was selected by minimising the Bayesian Information Criterion (BIC) over *k ∈* {1, *…*, 6} ; the optimal *k* was identified as the elbow of the BIC curve. Individual Gaussian components are shown as coloured curves, with the mixture density overlaid (dashed black). Each component is annotated with its mean *µ*, standard deviation *σ*, and mixture weight.

#### D.5 Additional Animals: Delay Distribution Across 100 Random Train-Test Splits

**Figure 12:**
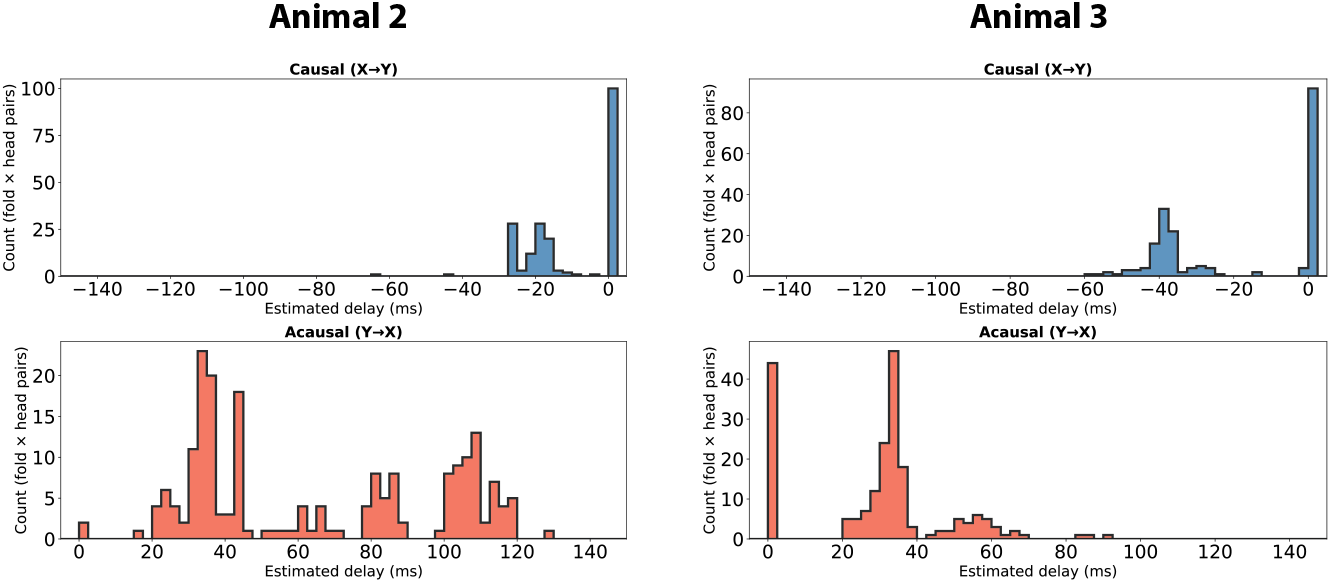
Estimated Delay Distributions for different animals. We performed the same random train-test split discussed in section 4.2 for two different sessions from two different animals and we found distinct characteristic lags across animals, reflecting differences in recording sites and the specific neural populations sampled within the IRN/PARN

#### D.6 Attention Heads and Diagonal Profiles for Multi-Region Neural Experiment

**Figure 13:**
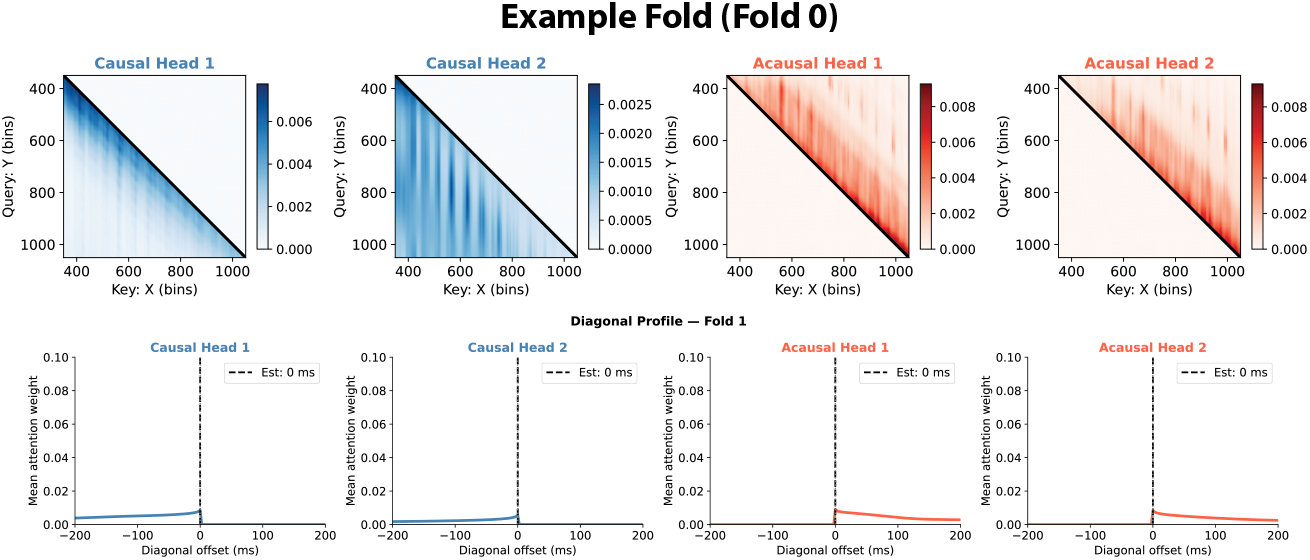
Zoomed-in attention maps (top) and diagonal profiles (bottom) for an example fold of the multi-region (S1–M1) experiment. All four heads — two causal and two acausal — show attention weights that decay monotonically from the main diagonal, with no peak at a non-zero offset, indicating that the model does not identify a dominant temporal lag between S1 and M1 activity in this session.

## Notes

### Competing Interest Statement

The authors have declared no competing interest.

## References

[1] Adrian G. Bondy et al. Brain-wide coordination of internal signals during decision-making. bioRxiv, 2024.

[2] Susu Chen et al. Brain-wide neural activity underlying memory-guided movement. Cell, 187(3):676–691, 2024.

[3] Nicholas A. Steinmetz, Christof Koch, Kenneth D. Harris, and Matteo Carandini. Challenges and opportunities for large-scale electrophysiology with Neuropixels probes. Current Opinion in Neurobiology, 50:92–100, 2018.

[4] Andrei Khilkevich et al. Brain-wide dynamics linking sensation to action during decision-making. Nature, 634(8035):890–900, 2024.

[5] James J. Jun et al. Fully integrated silicon probes for high-density recording of neural activity. Nature, 551(7679):232–236, 2017.

[6] Earl K. Miller, Mikael Lundqvist, and Andre M. Bastos. Working Memory 2.0. Neuron, 100:463–475, 2018.

[7] Munib A. Hasnain, Jaclyn E Birnbaum, et al. Separating cognitive and motor processes in the behaving mouse. Nature Neuroscience, 28:640–653, 2025.

[8] Victor A. Lamme, Hans Supèr, and Henk Spekreijse. Feedforward, horizontal, and feedback processing in the visual cortex. Current Opinion in Neurobiology, 8:529–535, 1998.

[9] Alessandra Angelucci and Paul C. Bressloff. Contribution of feedforward, lateral and feedback connections to the classical receptive field center and extra-classical receptive field surround of primate V1 neurons. Progress in Brain Research, 154:93–120, 2006.

[10] Charles D. Gilbert and Wu Li. Top-down influences on visual processing. Nature Reviews Neuroscience, 14:350–363, 2013.

[11] Kenneth D. Harris and Thomas D. Mrsic-Flogel. Cortical connectivity and sensory coding. Nature, 503:51–58, 2013.

[12] Reza Shadmehr and John W. Krakauer. A computational neuroanatomy for motor control. Experimental Brain Research, 185:359–381, 2008.

[13] Johannes M. Mayrhofer et al. Distinct contributions of whisker sensory cortex and tongue-jaw motor cortex in a goal-directed sensorimotor transformation. Neuron, 103:1034–1043, 2019.

[14] Da Xu et al. Cortical processing of flexible and context-dependent sensorimotor sequences. Nature, 603:464–469, 2022.

[15] Tudor Dragoi et al. Dynamic engagement of the motor cortex in controlling movement. bioRxiv, 2026.

[16] Sander W. Keemink and Christian K. Machens. Decoding and encoding (de)mixed population responses. Current Opinion in Neurobiology, 58:112–121, 2019.

[17] Matthew T. Schmolesky et al. Signal timing across the macaque visual system. Journal of Neurophysiology, 79:3272–3278, 1998.

[18] Joshua Glaser, Matthew Whiteway, John P. Cunningham, Liam Paninski, and Scott Linderman. Recurrent switching dynamical systems models for multiple interacting neural populations. Advances in Neural Information Processing Systems, 33:14867–14878, 2020.

[19] B. Liu, J. Sacks, and Matthew D. Golub. Accurate identification of communication between multiple interacting neural populations. arXiv, 2025.

[20] O. Karniol-Tambour et al. Modeling state-dependent communication between brain regions with switching nonlinear dynamical systems. In The Twelfth International Conference on Learning Representations, 2023.

[21] Matthew G. Perich and Kanaka Rajan. Rethinking brain-wide interactions through multi-region ‘network of networks’ models. Current Opinion in Neurobiology, 65:146–151, 2020.

[22] Evren Gokcen et al. Disentangling the flow of signals between populations of neurons. Nature Computational Science, 2:512–525, 2022.

[23] Joao D. Semedo, Amin Zandvakili, Christian K. Machens, Byron M. Yu, and Adam Kohn. Cortical areas interact through a communication subspace. Neuron, 102:249–259, 2019.

[24] Byungwoo Kang and Shaul Druckmann. Approaches to inferring multi-regional interactions from simultaneous population recordings. Current Opinion in Neurobiology, 65:108–119, 2020.

[25] Stephen L. Keeley, David M. Zoltowski, Mikio C. Aoi, and Jonathan W. Pillow. Modeling statistical dependencies in multi-region spike train data. Current Opinion in Neurobiology, 65:194–202, 2020.

[26] Evren Gokcen et al. Uncovering motifs of concurrent signaling across multiple neuronal populations. Advances in Neural Information Processing Systems, 2023.

[27] Saurabh Vyas, Matthew D. Golub, David Sussillo, and Krishna V. Shenoy. Computation through neural population dynamics. Annual Review of Neuroscience, 43:249–275, 2020.

[28] Scott W. Linderman et al. Recurrent switching linear dynamical systems. arXiv, 2016.

[29] Chethan Pandarinath et al. Inferring single-trial neural population dynamics using sequential auto-encoders. Nature Methods, 15:805–815, 2018.

[30] Amber Hu et al. Modeling latent neural dynamics with Gaussian process switching linear dynamical systems. arXiv, 2024.

[31] Noga Mudrik, Yenho Chen, Eva Yezerets, Christopher J. Rozell, and Adam S. Charles. Decom-posed linear dynamical systems (dLDS) for learning the latent components of neural dynamics. Journal of Machine Learning Research, 25:1–44, 2024.

[32] Valerio Mante, David Sussillo, Krishna V. Shenoy, and William T. Newsome. Context-dependent computation by recurrent dynamics in prefrontal cortex. Nature, 503:78–84, 2013.

[33] Laura N. Driscoll et al. Dynamic reorganization of neuronal activity patterns in parietal cortex. Cell, 170:986–999, 2017.

[34] Harold Hotelling. Relations between two sets of variates. Biometrika, 28:321–377, 1936.

[35] Joel Ye, Jennifer L. Collinger, Leila Wehbe, and Robert Gaunt. Neural data transformer 2: Multi-context pretraining for neural spiking activity. bioRxiv, 2023.

[36] Mehdi Azabou et al. A unified, scalable framework for neural population decoding. arXiv, 2023.

[37] Mehdi Azabou et al. Multi-session, multi-task neural decoding from distinct cell-types and brain regions. In International Conference on Learning Representations, 2025.

[38] Yizi Zhang et al. Neural encoding and decoding at scale. arXiv, 2025.

[39] Ram Dyuthi Sristi et al. Coupled transformer autoencoder for disentangling multi-region neural latent dynamics. arXiv, 2025.

[40] Qi Xin and Robert E. Kass. Identifying interactions across brain areas while accounting for individual-neuron dynamics with a transformer-based variational autoencoder. arXiv, 2025.

[41] Jianlin Su et al. Roformer: Enhanced transformer with rotary position embedding. Neurocomputing, 568:127063, 2023.

[42] Alan Edelman, Tomás A. Arias, and Steven T. Smith. The Geometry of Algorithms with Orthogonality Constraints, volume 20. 1998.

[43] Jun Takatoh et al. Constructing an adult orofacial premotor atlas in Allen mouse CCF. eLife, 10:e67291, 2021.

[44] Heet Kaku, Liu D. Liu, Runbo Gao, Steven West, Song-Mao Liao, Arseny Finkelstein, David Kleinfeld, Alyse Thomas, Sri Laasya Tipparaju, Karel Svoboda, and Nuo Li. A brainstem map of orofacial rhythms. bioRxiv, page 2025.01.27.635041, 2025.

[45] Diederik P Kingma and Jimmy Ba. Adam: A method for stochastic optimization. International Conference on Learning Representations (ICLR), 2015.

